# Nicotine dependence and functional connectivity of insular cortex subregions

**DOI:** 10.1101/2021.07.07.451360

**Authors:** Dara G. Ghahremani, Jean-Baptiste Pochon, Maylen Perez Diaz, Rachel F. Tyndale, Andy C. Dean, Edythe D. London

**Affiliations:** Department of Psychiatry and Biobehavioral Sciences, Semel Institute for Neuroscience and Human Behavior, University of California, Los Angeles, CA, USA; Department of Pharmacology & Toxicology and Department of Psychiatry, University of Toronto, 1 King’s College Circle, Toronto, ON, Canada; Campbell Family Mental Health Research Institute, Centre for Addiction & Mental Health, Toronto, ON, Canada; Department of Molecular and Medical Pharmacology, and University of California, Los Angeles, CA, USA; Brain Research Institute, University of California, Los Angeles, CA, USA

**Author notes:** Contributed equally to this work (Dara G. Ghahremani, Jean-Baptiste Pochon). Corresponding Authors: Dara Ghahremani, Ph.D. & Edythe D. London, Ph.D. Semel Institute for Neuroscience and Human Behavior, 760 Westwood Plaza, Room C8-831, Los Angeles, CA 90095-1759.

**Keywords:** Nicotine dependence, fMRI, insula, resting state functional connectivity

## Abstract

The insular cortex has been identified as a promising target in brain-based therapies for Tobacco Use Disorder, and has three major sub-regions (ventral anterior, dorsal anterior, and posterior) that serve distinct functional networks. How these subregions and associated networks contribute to nicotine dependence has not been well understood, and therefore was the subject of this study. Forty-seven individuals (24 women; 18-45 years old) who smoked cigarettes daily rated their dependence using the Fagerström Test for Nicotine Dependence (FTND), abstained from smoking overnight (~12 h), and underwent resting-state functional MRI. Correlations between dependence and resting-state functional connectivity (RSFC) of the major insular sub-regions were evaluated using whole-brain-corrected voxel-wise analyses and post-hoc region-of-interest (ROI) analyses. Dependence was analyzed both as a unitary (FTND total score) and bivariate construct – two FTND factors (“morning smoking” and “daytime smoking”). Dependence was negatively correlated with connectivity of both the right dorsal and left ventral anterior insula with the left precuneus, and with connectivity of the left posterior insula to the left putamen. In post-hoc analyses, dependence correlated negatively with connectivity between all anterior insula subregions and the left precuneus, and with bilateral posterior insula connectivity with the left posterior putamen. The latter finding was driven by “daytime smoking”. These results suggest an anterior-posterior distinction in functional insular networks associated with different dimensions of nicotine dependence, with greater dependence linked to weaker connectivity. They may inform therapeutic approaches involving brain stimulation that may elicit differential clinical outcomes depending on the insular subnetwork targeted.

## Introduction

The use of combustible tobacco products continues to be a substantial global health problem, causing over 7 million or 1 in 10 deaths world-wide each year ^1, 2^. Among people who try to quit smoking independently, only 3 to 6 percent successfully stop for 6 to 12 months, with most failing within 8 days ^3^. Behavioral and pharmacological treatments for Tobacco Use Disorder (TUD) also have limited success ^4–7^, and the need for investigation of novel therapeutic strategies persists. Elucidating the neural mechanisms of nicotine dependence has the potential to advance the treatment of TUD ^8, 9^, particularly with brain-based approaches, such as targeted brain stimulation ^10^. Knowledge of the neural systems underlying nicotine dependence is critical for guiding these treatments.

The insula has been identified as a promising target for brain-based treatments for TUD ^11, 12^, partly due to clinical evidence for its role in smoking behavior. Patients with insula lesions resulting from strokes have exhibited marked reduction in smoking ^13–16^. In stroke survivors, damage to the right insula resulted in smoking cessation when assessed one year after discharge from the hospital ^16^, and predicted even greater reduction in dependence when combined with damage to the basal ganglia ^14^.

Neuroimaging studies have also indicated the importance of the insula in maintenance of cigarette smoking. Cortical thickness of insular sub-regions is negatively related to nicotine dependence ^17–19^ and cigarette craving ^20^. Resting-state functional connectivity (RSFC) has been used to assess neural systems involved in nicotine dependence ^21, 22^. Studies that focused on the anterior cingulate cortex (ACC) found a negative relationship between nicotine dependence and connectivity with the striatum ^23–26^, and a rodent study showed that insula-frontal connectivity moderates this negative association ^27^. Other RSFC studies showed an inverse relationship between nicotine dependence and insula-ACC connectivity in humans ^24, 28, 29^.

The insula has been subdivided into three sub-regions that serve distinct functions: dorsal anterior, ventral anterior, and posterior ^30–32^. While the posterior portion of the insula connects with sensorimotor integration areas (e.g., pre-motor, supplementary motor cortex), the anterior portion is functionally linked with limbic regions and is a key component of the “salience network”, which includes the ACC ^33, 34^. The anterior portion has generally been shown to serve cognitive and affective functions ^35, 36^. Further functional distinctions have been made between dorsal and ventral anterior insula connectivity along cognitive and affective domains, respectively ^35, 37, 38^. In addition, dorsal/ventral distinctions have been conceptualized as components of externally- and internally-oriented networks – specifically, the frontoparietal attention network and the default mode network, respectively ^39, 40^.

Prior studies of RSFC have attempted to distinguish between connectivity of insular sub-regions and smoking-related behavioral variables ^24, 40–43^, but only two of them examined RSFC of insula subregions with respect to nicotine dependence ^24, 28^. In one of these studies, analysis was restricted to the dorsal ACC after demonstration of a difference in anterior insula-ACC connectivity between people who smoked and those who did not; a negative correlation between dependence and anterior insula connectivity with the dorsal ACC was found ^28^. In the other study, individuals who smoked and had schizophrenia were compared with individuals who smoked but had no other psychiatric diagnosis ^24^. Both groups showed an inverse relationship between nicotine dependence and posterior insula-dorsal ACC connectivity. Despite this initial evidence for distinctions between circuits of insula subregions with respect to dependence, a comprehensive analysis of the contributions of the three major subregions is lacking, leaving open the possibility that RSFC of different insular subregions would be differentially related to nicotine dependence. We therefore undertook a comprehensive analysis of correlations with connectivity patterns of these finer-grained subregions in a whole-brain analysis.

The Fagerström Test for Nicotine Dependence (FTND) ^44^ is a widely used, validated assessment that is related to biological indices of smoking ^45, 46^. Neuroimaging studies that evaluate nicotine dependence with the FTND have used a unidimensional scoring system; however, psychometric studies have established that a two-dimensional (two factor) structure is more appropriate ^47–51^, with the first factor interpreted as “the degree of urgency to restore nicotine levels to a given threshold after nighttime abstinence” (“morning smoking”) and a second factor interpreted as “the persistence with which nicotine levels are maintained at a given threshold during waking hours” (“daytime smoking”) ^51^. To date, no neuroimaging studies have examined the neural circuitry underlying this bivariate structure of the FTND, which may provide a more nuanced understanding of nicotine dependence and the neural circuits that support it.

In a group of 47 participants who smoked cigarettes daily, we examined the relationship between nicotine dependence and RSFC of the three major insular sub-regions (ventral anterior, dorsal anterior, and posterior) after overnight abstinence. Considering the literature, we hypothesized that dependence would be negatively correlated both with connectivity between anterior insula subregions and ACC, and with connectivity between the posterior insula and sensory-motor integration regions (e.g., supplementary motor area) (i.e., participants with greater connectivity would show less dependence). To determine the extent to which the two-factor structure reveals greater specificity with respect to the relationship between dependence and insula connectivity, we examined insula RSFC in relation to both the unidimensional scoring (FTND total score) and the two-factors.

## Materials and Methods

### Overview of Experimental Design

Functional magnetic resonance imaging (fMRI) data were collected during the resting state from adults who smoked cigarettes daily and maintained overnight abstinence before testing. The study was part of a larger investigation on the brain correlates of smoking behavior and took place between September, 2017 and February, 2020. Another report from that study is in press ^52^. The study was conducted at the Semel Institute for Neuroscience and Human Behavior at the University of California, Los Angeles (UCLA). All study procedures were approved by the UCLA Institutional Review Board.

### Participants

One-hundred-seventy-nine participants were screened via online and print advertisements. They attended an intake session where they received a detailed explanation of the study procedures, provided written informed consent, and were screened for eligibility. Fifty-one met all study criteria and completed all procedures. Inclusion criteria were as follows: age of 18-45 years, generally good health, self-report of smoking at least 4 cigarettes per day for at least 1 year, and urinary cotinine ≥100 ng/mL. Recent smoking history was verified during the intake session using a urine cotinine test (ACCUTEST Urine Cotinine Test, Jant Pharmacal Corp., Encino, CA, score ≥3, cotinine ≥100 ng/mL). Exclusion criteria were positive urine tests for drugs of abuse other than nicotine or tetrahydrocannabinol, consuming ≥10 alcoholic drinks per week, any current psychiatric disorder other than Tobacco Use Disorder as assessed via the Mini International Neuropsychiatric Interview (MINI) for DSM-5^53, 54^, history of neurological injury, and using electronic cigarettes, cigars, snuff, or chewing tobacco >3 times a month.

### Verification of Drug and Alcohol Abstinence

On the testing day, overnight (~12 hours) abstinence from smoking was verified by a CO level of <10 ppm measured with the Micro+ Smokerlyzer® breath CO monitor (Bedford Scientific Ltd., Maidstone, Kent, UK). Abstinence from cocaine, opiates, benzodiazepines, and amphetamines was also verified with a five-panel urine drug test (Drugs of Abuse Test Insta-view®, Alfa Scientific Designs Inc., Poway, CA). Alcohol abstinence was verified using a breathalyzer (Alco-Sensor FST®, Intoximeters, Inc., St. Louis, MO). Recent abstinence from cannabis use was verified with the Dräger DrugTest® 5000 saliva test (Dräger, Inc., Houston, TX).

### Self-report of Nicotine Dependence and Analysis Procedures

Nicotine dependence was measured during the intake session using FTND ^44^. The standard total score was computed in addition to the two-factor solution. These factors have been established in the literature using exploratory factor analyses ^47–50^ and confirmatory factor analysis ^51^, indicating that the six items of the FTND comprise two factors with one item (time to first cigarette of the day) loading on both factors. The first factor included items 1, 3 and 5 of the FTND (FTND135) and is interpreted as encompassing “morning smoking”, and the second factor is composed of items 1, 2, 4 and 6 (FTND1246), and is interpreted as “daytime smoking” (see Supplementary Table 13 for FTND items). For each participant, we computed a weighted sum of the response to the items in each factor, with weightings based on the mean of the weights reported across previous studies ^47–50^. See supplementary materials for analysis details.

We verified that FTND data from our 47 participants matched the two-factor structure determined in the literature on much larger samples. Rather than conducting a confirmatory factorial analysis that requires a large number of observations, we performed the same factorial analysis method conducted in the study by Haddock et al. ^48^ on 4,042 smokers which used principal component analysis followed by a Varimax rotation. Results of this decomposition (Supplementary Table 13) showed that our data exhibited the same two-factor organization. Namely, FTND135 and FTND1246 explained 54% of the variance in our study and 51% in the study of Haddock et al. The average difference in item loading for FTND135 and FTND1246 between both studies was 0.14 and 0.06, respectively.

### Scanning Protocol

#### MRI data acquisition

All images were acquired on a 3-Tesla PRISMA (Siemens) MRI scanner using a 32-channel head coil receiver. The resting state imaging protocol consisted of the continuous acquisition of 738 Echo-planar Image (EPI) volumes over a period of 9 minutes and 50 seconds. A multi-band accelerated EPI pulse sequence (factor 8) was used, allowing us to acquire 72 axial slices during a repetition time (TR) of 800 ms with a 104×104 matrix. Resolution was 2×2×2 mm^3^, echo time (TE) was 37 ms, and the flip angle was 52 degrees. Participants were asked to keep their eyes open and to look at a black screen during the resting state scan. The structural T1-weighted images were obtained using a Magnetization Prepared Rapid Gradient Echo (MPRAGE) sequence with the following parameters: isovoxel 0.8 mm^3^, FOV = 240 × 256 mm^2^, TE = 2.24 ms, TR = 2400 ms; flip angle = 8°; 208 sagittal slices.

#### MRI data pre-processing

Image preprocessing was mostly conducted with FSL (5.0.9). The initial stages included rigid body realignment to correct for head movements within each scanning run, skull removal, and non-linear registration to the Montreal Neurological Institute (MNI) template. A first motion cleaning and noise reduction were performed using a 24-parameter linear regression model that included six motion parameters (3 translational dimensions along X, Y and Z axes and 3 rotational dimensions: “pitch”, “roll” and “yaw”), the temporal derivatives of these parameters and the quadratic of all parameters ^55^. Mean frame displacement (FD) and the variance of signal change from the average signal (DVARS) of the raw images were estimated. A null sampling distribution of DVARS was used to identify frames with excessive variance at p < 0.05 ^56^; frames with FD exceeding 0.45 mm were also flagged. These frames as well as the one located in time just prior (t-1) and two just after (t+1 and t+2) were included in a censoring temporal mask for data interpolation: a least-squares spectral decomposition of the uncensored data was performed to reconstitute data of the censored timepoints see methods in ^57^. The uncensored data defined the frequency characteristics of signals that then replaced the censored data. This step aimed at minimizing the contamination of the signal from the censored frames during frequency filtering. The interpolated signal was then demeaned, detrended and filtered using an ideal bandpass filter (0.009 – 0.08 Hz). Following band-pass filtering, the interpolated timepoints were finally censored. Participants with more than 50% frames censored (i.e., those with less than 5 minutes of remaining resting state data) were excluded from the analysis. To reduce the contribution from non-neuronal noise in the data, the minimal number of principle components that explained at least 50% of the variance of mean signal extracted from white matter and cerebrospinal fluid were evaluated and regressed out from the signal aCompCor50, ^58^. Volumes were then spatially smoothed with a Gaussian filter using a 5-mm FWHM kernel. Each voxel was normalized to a mean value of 100 (SD=1) to transform the data to Pearson’s correlation coefficients (r). All analyses were performed on Linux (CentOS release 6.10) using FSL 5.0.9, MATLAB 8.6, R (version 3.6.0).

#### Resting-state fMRI seed-based analysis

To minimize bias, we used a statistically conservative voxel-wise whole-brain analytic approach rather than restricting to a priori-selected target regions or networks. On each of the two hemispheres of the brain three insula seeds, which encompassed ventral-anterior, dorsal-anterior, and posterior aspects of the insula, were defined for RSFC analyses (Figure 1). To define the anterior insula, we compared anatomical landmarks from a probabilistic atlas ^59^ to functional connectivity-based parcellations of the insula ^30, 32^. From these studies, we defined the ventral anterior insula parcel as the anterior inferior insular cortex (which includes the apex, the limen, and the transverse gyrus). The dorsal anterior insula was defined as the anterior and middle short gyri. The precentral sulcus was used to segment the anterior from the posterior insula. Using these landmarks, we manually determined the anterior insular subdivisions (dorsal and ventral, left and right) from the MNI152 template.

**Figure 1.**
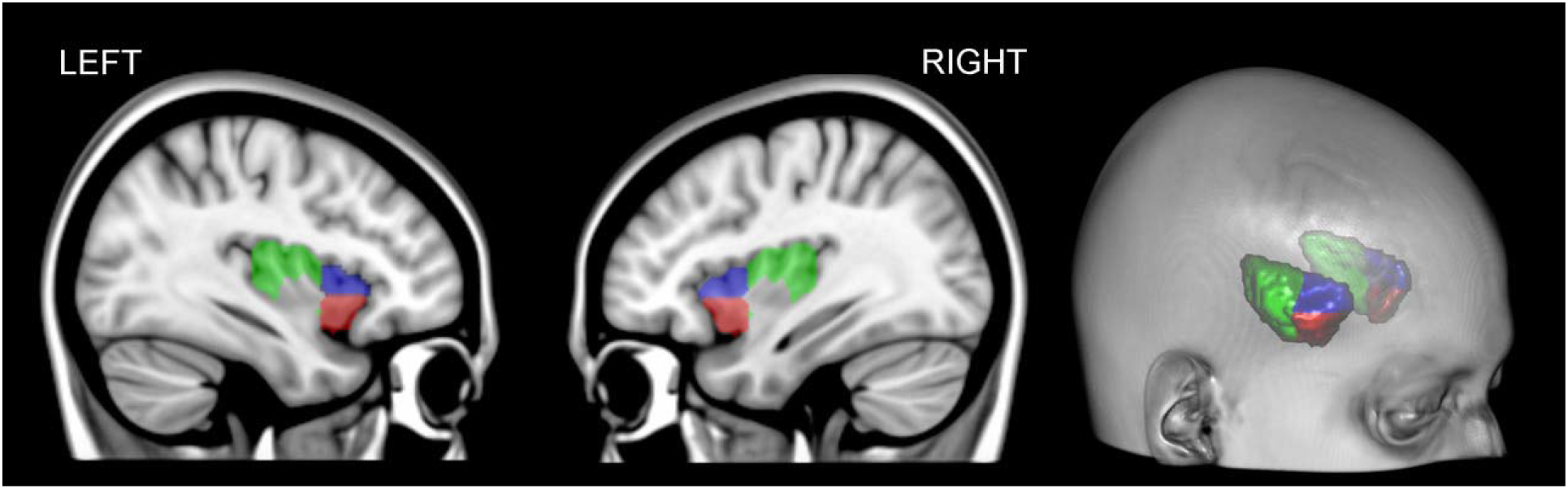
Insula sub-regions used as seeds for resting state functional connectivity analyses. Red – ventral anterior insula, Blue – dorsal anterior insula, Green – posterior Insula.

To evaluate the functional connectivity between the insula seeds and other brain regions during the resting state, the time series from each seed was extracted, and its first normalized eigen vector (mean=100, SD=1; to facilitate computation of Pearson’s r) was used as a regressor in an ordinary least squares linear regression analysis on every voxel (as implemented in film_gls in FEAT). The parameter estimates of the model, corresponding to the Pearson’s correlation coefficient (since data were previously normalized), were z-transformed to improve data normality.

The resulting z-transformed images were used in multi-level mixed effects models for group analyses (FLAME1, FEAT) testing for the effect of nicotine dependence on functional connectivity for each seed. Specifically, two separate models were tested. The first model included the total score of the FTND as the independent variable of interest. The second model examined separate effects of the two factors of the FTND: FTND135 (“morning smoking”) and FTND1246 (“daytime smoking”).

To account for differences in motion during scanning between participants, the mean frame displacement value was included as a covariate in all models in addition to age. Results were cluster-corrected for multiple comparisons using a voxel-height threshold of p < 0.001 (Z>3.1) and cluster threshold of p<0.05 as recommended per Eklund et al ^60^. The coordinates reported here correspond to the peak voxel within a given cluster in MNI coordinate space. All post-hoc tests on data extracted from statistical maps were conducted using R ^61^. For each participant, data were extracted and averaged from voxels within clusters that survived the cluster-correction threshold (voxel height = Z>3.1, cluster extent = P<0.05).

## Results

### Participant Characteristics

Fifty-one adults who endorsed daily cigarette smoking completed overnight (~12 hours) abstinence from smoking. Of the 51 participants tested, four were excluded for excessive motion during fMRI, as revealed by the number of flagged volumes exceeding DVARS and/or FD thresholds after data preprocessing (>50%). The final sample included 47 individuals (24 female) with a mean age of 33.3 (SD=7.2) years. On average, they smoked 11.4 (SD=4.5) cigarettes per day with a smoking history of 8.1 (SD=6.0) pack years. Nicotine dependence varied from low to high levels (mean FTND total scores of 4.0, SD=2.0). Other participant characteristics, including ethnicity, race, alcohol, and cannabis consumption, are shown in Table 1.

**Table 1.** Participant Characteristics (n=47)

|  |  |  |
| --- | --- | --- |
| Sex (F/M) | n=24 / n=23 |  |
| Ethnicity |  |  |
| Hispanic/Latino (12.8%) |  |  |
| Non-Hispanic/Latino (83%) |  |  |
| Unknown (4.3%) |  |  |
| Race |  |  |
| African-American (23.4%) |  |  |
| Asian (2.1%) |  |  |
| Caucasian (59.6%) |  |  |
| Mixed (8.5%) |  |  |
| Pacific Islander (4.3%) |  |  |
| Unknown (2.1%) |  |  |
|  | Mean | (SD) |
| Age (years) | 33.3 | (7.2) |
| Nicotine Dependence<sup>a</sup> | 4.0 | (2.0) |
| Tobacco / current use, |  |  |
| cigarettes/day | 11.4 | (4.5) |
| Tobacco / lifetime exposure, |  |  |
| pack-years | 8.1 | (6.0) |
| Alcohol consumption (drinks/week) |  |  |
| n=24 (>1 drink / week) | 4.3 | (1.99) |
| Cannabis consumption (grams/week), |  |  |
| n=13 (>1 gram / week) | 3.92 | (5.19) |

<sup>a</sup>Fagerström Test of Nicotine Dependence

### Nicotine dependence

Testing the association of the FTND total score with functional connectivity between the insula seeds and the rest of the brain revealed negative relationships in all cases. Specifically, FTND was negatively correlated with: (1) connectivity of the left ventral anterior insula (lvaInsula) and the precuneus (X=−12, Y=−70, Z=48; Figure 2A, B), (2) connectivity of the right dorsal anterior insula (rdaInsula) and a precuneus target with very similar coordinates (X=−10, Y=−72; Z=50) (Figure 2A, B, C), and (3) connectivity between the left posterior insula (lpInsula) seed and the left posterior aspect of the putamen (X=−28, Y=−12, Z=2) (Figure 3A, C, D).

**Figure 2.**
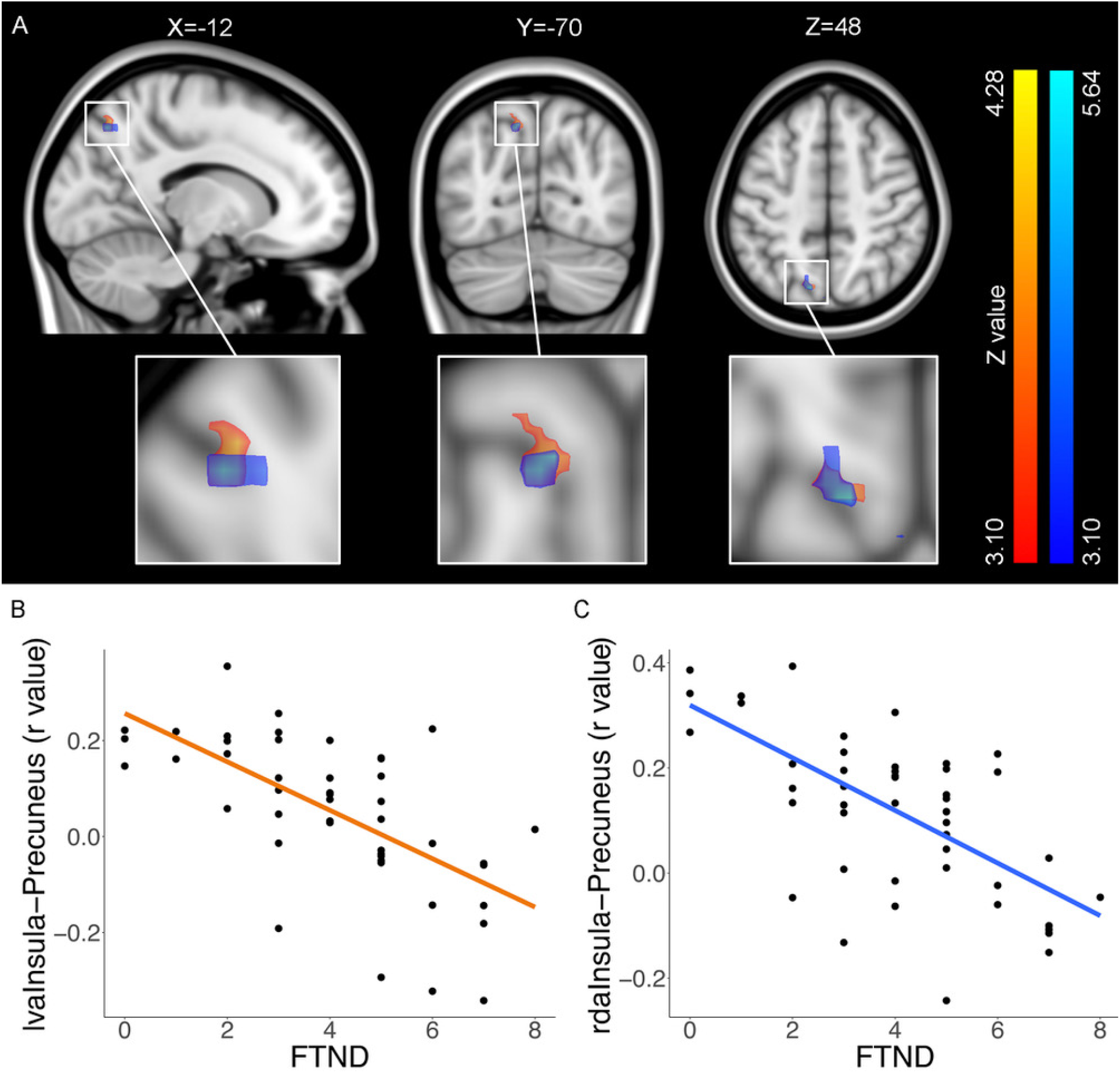
Relationship between nicotine dependence (FTND total score) with anterior insula-precuneus connectivity. **A**. Thresholded statistical maps indicating functional connectivity of left ventral anterior (hot colors) and right ventral dorsal (cool colors) with a cluster in the left precuneus correlating with FTND total score (voxel threshold: Z>3.1, cluster threshold: P<0.05). Brain images are presented in neurological convention (right=right). **B**. Scatterplots of data extracted from the left precuneus cluster from individual participants are shown along with linear fits to illustrate the negative direction of the relationship between FTND and functional connectivity.

**Figure 3.**
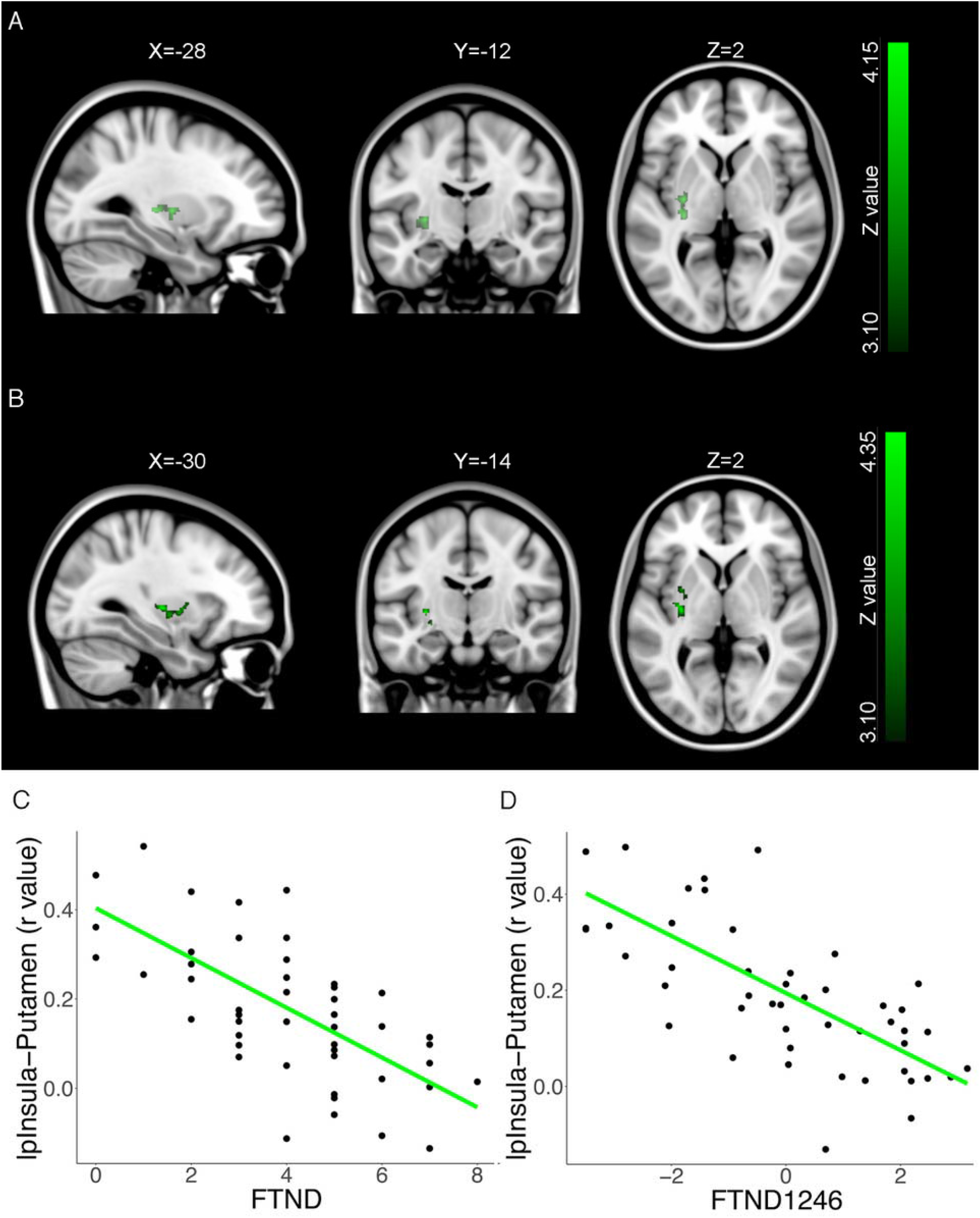
Relationship between nicotine dependence with posterior insula-left putamen connectivity. Thresholded statistical maps indicating functional connectivity of left posterior insula with a cluster in the left posterior putamen correlating with (**A**) FTND total score and (**B**) the “daytime smoking” factor (FTND1246) from the two-factor model of FTND (voxel threshold: Z>3.1, cluster threshold: P<0.05). **C.** Scatterplots of data extracted from the left putamen clusters from individual participants are shown along with linear fits to illustrate the negative direction of the relationship between the FTND measures and functional connectivity.

The second model, which tested for separate effects of the two FTND factors, revealed a negative relationship between “daytime smoking” (FTND1246) and connectivity of the lpInsula and the same left posterior putamen area found in analysis of the FTND total score (X=−30, Y=− 14; Z=2; Figure 3B, C, D). No other significant relationships were observed, including any with the “morning smoking” factor (FTND135).

### Post hoc analyses of associations with nicotine dependence

The results presented above were determined in whole-brain voxel-wise analyses that involved a conservative correction for multiple comparisons. We noted the asymmetry of the insular seeds that yielded the patterns observed and aimed to explore the extent of this asymmetry using post-hoc region-of-interest analyses. Accordingly, we conducted region-of-interest analyses testing for relationships between dependence and connectivity between all four anterior insula seeds (tested separately) and the left precuneus cluster as well as the two posterior insula seeds and the left putamen cluster. These analyses were conducted using the same two linear models – one with FTND total as independent variable and another with FTND135 and FTND1246. The results indicated that functional connectivity of all four anterior insula seeds (not only the rdaInsula and lvaInsula) with the precuneus was negatively correlated with the total FTND score at *p*s< 0.001 (Supplementary Tables 1-4) and that both left and right posterior insula connectivity with the left posterior putamen had the same modulatory pattern (*p*s < 10^−5^) (Supplementary Tables 5 and 6).

The test of the two-factor model indicated that, similar to findings for lpInsula, rpInsula connectivity with the left posterior putamen was also modulated by the “daytime smoking” factor (FTND1246) (*p*<2×10^−6^) (Supplementary Tables 7 and 8). Connectivity of all anterior seeds of the insula with the precuneus showed significant negative relationships with both factors, except for the rvaInsula (p=0.08) (Supplementary Tables 9-12). Table 2 summarizes these *post hoc* results.

**Table 2.** Summary of post-hoc region of interest analyses of the right precuneus and left putamen for testing contra-lateral insula seeds. Results of each post-hoc linear model is presented in Supplementary Tables 1-12.

| Insula seed | Right precuneus |  |  |
| --- | --- | --- | --- |
|  | FTND<br>Total | “Morning<br>Smoking”<sup>a</sup> | “Daytime<br>Smoking”<sup>b</sup> |
| left ventral anterior insula | *** | * | ** |
| left dorsal anterior insula | *** | * | * |
| right ventral anterior insula | *** | * |  |
| right dorsal anterior insula | *** | ** | * |

| Insula seed | Left putamen |  |  |
| --- | --- | --- | --- |
|  | FTND<br>Total | “Morning<br>Smoking”<sup>a</sup> | “Morning<br>Smoking”<sup>a</sup> |
| left posterior insula | *** |  | *** |
| right posterior insula | *** |  | *** |

\* = $p < 0.05$ , \*\* = $P < 0.01$ , \*\*\* = $P < 0.001$
<sup>a</sup> “Morning Smoking” was associated with FTND items 1, 3, and 5
<sup>b</sup> “Daytime Smoking” was associated with FTND items 1, 2, 4, and 6

## Discussion

This study showed that nicotine dependence was negatively correlated with two distinct functional connectivity patterns across different insular subregions. Taken together, the combined results of *a priori* whole-brain and post-hoc region-of-interest analyses indicated that nicotine dependence is associated with connectivity of the anterior insula and the right precuneus and with connectivity of the posterior insula and the left posterior putamen. Moreover, when examining two separate aspects of dependence (as defined by the two-factor model), anterior insula-right precuneus connectivity was related to both “morning smoking” and “daytime smoking” whereas posterior insula-left putamen connectivity was only related to “daytime smoking.” Thus, different dimensions of dependence apparently are related to connectivity of separate insular subregions.

This separation of anterior vs. posterior insula functional connectivity is somewhat consistent with functional distinctions observed along the anterior/posterior insula axis (affective-cognitive/sensorimotor, respectively) ^35^. Although we hypothesized a relationship between dependence and connectivity of anterior insula subregions with limbic regions and components of the salience network (i.e. ACC), we found a relationship with connectivity to the precuneus, a region that is not included among limbic or salience network structures but is involved in cognitive and affective function. The precuneus has been associated with self-referential processing, empathy, and episodic memory retrieval ^62^, and has a central role in the default mode network ^63–65^. In studies of substance use, the precuneus is among the primary brain regions activated during smoking cue-induced neural reactivity, as indicated by meta-analyses ^66^. Precuneus activation during presentation of smoking- and alcohol-related cues is positively associated with nicotine and alcohol dependence, respectively ^67^. Moreover, in individuals who smoke cigarettes, anterior insula-precuneus connectivity, a similar connectivity pattern to that observed in the current study, was correlated with cue-induced craving ^68^. Combined with these findings from prior studies, those presented here suggest a role for the interaction of the precuneus with the anterior insula in maintenance of nicotine dependence, especially aspects that involve self-referential processes.

The relationship between nicotine dependence and posterior insula-left posterior putamen connectivity is consistent with the view that the insula supports an embodied experience of addiction via integration of interoceptive signals ^69^, in light of the role of the posterior insula in interoception ^70–72^ and evidence for a role of the putamen in habitual stimulus-response associations ^73^. A study of stroke patients comparing those with basal ganglia lesions (including the putamen), and those with both insula and basal ganglia lesions, showed that patients with both lesions had a greater disruption of smoking after their stroke than patients with basal ganglia lesions alone ^14^, providing evidence for the relevance of insula-basal ganglia interactions in maintaining dependence. Our functional connectivity finding is also supported by a structural connectivity study that mapped insular subregions to subcortical regions, indicating that of the subcortical regions tested (thalamus, amygdala, hippocampus, putamen, globus pallidus, caudate nucleus, and nucleus accumbens), the putamen had the most connections with the posterior insula ^74^. Association of weaker posterior insula-left putamen functional connectivity with greater nicotine dependence is in line with meta-analytic findings of less engagement of striatal regions in response to smoking-related and non-smoking reward-related cues with greater nicotine dependence ^75^. The latter findings suggest that weak posterior insula-putamen connectivity may also be related to disrupted reward-related responses in the striatum.

This study is the first to examine neuroimaging markers of nicotine dependence with respect to the widely established two-factor characterization of dependence measured using the FTND. That posterior insula-left putamen connectivity uniquely correlated with the “daytime smoking” factor, and not “morning smoking”, suggests that this functional circuit especially serves persistence in maintaining nicotine levels throughout the day. Given that some of the items included in this factor may be considered as assessments of self-control (e.g., “Do you find it difficult to keep from smoking in places where it is not allowed?”), weakened connectivity of this circuit may lead to disruption of interoceptive signals that support self-control behavior. The relationship between anterior insula-precuneus connectivity and both factors suggests that this functional circuit is important for multiple dimensions of dependence.

Our results are only partially consistent with previous insula RSFC studies of nicotine dependence, which focused on the ACC and found significant associations involving the ACC. A previous study of non-deprived young participants (15-24 years of age) found a negative relationship between anterior insula-ACC connectivity and nicotine dependence ^28^; but ACC was selected as the connectivity target post-hoc, based on an analysis that compared anterior insula connectivity of individuals who did or did not smoke and found greater connectivity with the ACC as a primary group difference. Another study similarly found a negative association between dependence and insula-ACC connectivity ^29^, but it used the whole, bilateral insula as a seed, eliminating the possibility of differentiating effects of insular subregions. Lastly, a third study found a negative relationship between nicotine dependence and connectivity between the posterior insula and ACC in individuals with schizophrenia and also those without psychiatric diagnoses ^24^. It is possible that the current study did not find significant results involving the ACC because we used a whole-brain voxel-wise approach, with strict multiple comparison correction, to determine which connectivity targets across the whole brain were associated with dependence. Another reason for the absence of a positive finding regarding the ACC may be our use of rigorous motion-cleaning approaches ^57, 76, 77^, which are considered essential for removing artifacts in the data that may lead to spurious correlations ^78^.

Brain stimulation has been considered as a promising therapy for smoking cessation ^79, 80^, and studies targeting the insula have demonstrated mixed success in affecting smoking-related variables, such as craving and withdrawal ^10–12^. By highlighting the relevance of two insular functional connectivity patterns, which are weaker in strength in those who have greater nicotine dependence, we provide additional relevant targets for future stimulation studies. For example, stimulation studies may not only choose to target anterior and posterior insula, but also the right precuneus and left posterior putamen, with the aim of increasing the strength of the relevant functional connectivity patterns identified in the current study.

The current study is limited in that it cannot determine whether weak insular connectivity is a cause or consequence of nicotine dependence although pre-clinical studies have suggested a potential causal role of insular connectivity on dependence. In a data-driven study of rats, Hsu et al. ^81^ showed that an insular-frontal network module was predictive of later nicotine dependence. However, it remains to be determined if a similar finding will emerge from longitudinal human studies, such as the Adolescent Brain Development Study (ABCD) study ^82^.

Overall, this study provides evidence for the association of nicotine dependence with weak functional connectivity of distinct insula subregions, namely, anterior insula-right precuneus and posterior insula-left putamen. This regional dissociation within the insula highlights the heterogeneity of the insula with respect to neural processes involved in maintenance of smoking and suggests multiple potential targets for brain-based therapies that address nicotine addiction.

## Acknowledgements

The authors thank Ms. Andrea Donis, Ms. Diana Paez, Ms. Citlaly Cahuantzi, Ms. Tinisha Sakhrani, and Mr. Hector Diaz, whose contribution to data collection helped make this work possible.

## Author Contributions

All authors were involved in designing the study and contributed to writing the manuscript. Drs. Ghahremani and Perez Diaz acquired the data. Drs. Pochon, Ghahremani, Perez Diaz and Dean were responsible for data analysis. Dr. Tyndale contributed to decisions regarding participant characterization. Drs. Ghahremani, Pochon, and London drafted the manuscript. As principal investigator of the study, Dr. London is accountable for all aspects of the work, including its accuracy and integrity. All authors have reviewed and approve the final version of this manuscript.

## Data Availability

All self-report, toxicology, and summary fMRI data discussed in this manuscript, as well as the code used for statistical analyses, are publicly available from the Open Science Framework web site under project title, “Nicotine dependence and functional connectivity of insular cortex subregions” (https://osf.io/24tkf/).

