## Supplementary Tables for "Nicotine dependence and functional connectivity of insular cortex subregions"

### SUPPLEMENTAL TABLES

**Table 1**

**lvaInsula-precuneus = 1 + FTND + age + mFD**

| | Estimate | Std.Error | t-value | Pr(>\|t\|) | |
|---|---|---|---|---|---|
| (Intercept) | 0.176 | 0.106 | 1.66 | 0.104 | |
| FTND | -0.051 | 0.010 | -5.36 | 3.05E-06 | *** |
| Age | 0.001 | 0.003 | 0.45 | 0.657 | |
| mFD | 0.259 | 0.303 | 0.85 | 0.398 | |

Residual standard error: 0.1278 on 43 degrees of freedom

Multiple R-squared: 0.4055, Adjusted R-squared: 0.364

F-statistic: 9.775 on 3 and 43 DF, p-value: 4.882e-05

**Table 2**

**rdaInsula-precuneus = 1 + FTND + age + mFD**

| | Estimate | Std.Error | t-value | Pr(>\|t\|) | |
|---|---|---|---|---|---|
| (Intercept) | 0.284 | 0.101 | 2.80 | 0.00759 | ** |
| FTND | -0.051 | 0.009 | -5.63 | 1.27E-06 | *** |
| Age | -0.001 | 0.002 | -0.36 | 0.72423 | |
| mFD | 0.407 | 0.290 | 1.41 | 0.16677 | |

Residual standard error: 0.122 on 43 degrees of freedom

Multiple R-squared: 0.4319, Adjusted R-squared: 0.3922

F-statistic: 10.9 on 3 and 43 DF, p-value: 1.891e-05

**Table 3**

**ldaInsula-precuneus = 1 + FTND + age + mFD**

| | Estimate | Std.Error | t-value | Pr(>\|t\|) | |
|---|---|---|---|---|---|
| (Intercept) | 0.429 | 0.123 | 3.50 | 0.00109 | ** |
| FTND | -0.054 | 0.011 | -4.94 | 1.24E-05 | *** |
| Age | -0.002 | 0.003 | -0.68 | 0.49998 | |
| mFD | 0.224 | 0.350 | 0.64 | 0.52598 | |

Residual standard error: 0.1476 on 43 degrees of freedom

Multiple R-squared: 0.3656, Adjusted R-squared: 0.3213

F-statistic: 8.26 on 3 and 43 DF, p-value: 0.0001881

**Table 4**

**rvaInsula-precuneus = 1 + FTND + age + mFD**

| | Estimate | Std.Error | t-value | Pr(>\|t\|) | |
|---|---|---|---|---|---|
| (Intercept) | 0.055 | 0.139 | 0.39 | 0.69575 | |
| FTND | -0.048 | 0.012 | -3.87 | 0.00036 | *** |
| Age | 0.002 | 0.003 | 0.65 | 0.52241 | |
| mFD | 0.771 | 0.399 | 1.93 | 0.05979 | . |

Residual standard error: 0.168 on 43 degrees of freedom
Multiple R-squared: 0.2998, Adjusted R-squared: 0.251
F-statistic: 6.137 on 3 and 43 DF, p-value: 0.001438

**Table 5**

**lpInsula-putamen= 1 + FTND + age + mFD**

| | Estimate | Std.Error | t-value | Pr(>\|t\|) | |
|---|---|---|---|---|---|
| (Intercept) | 0.408 | 0.109 | 3.76 | 0.000507 | *** |
| FTND | -0.056 | 0.010 | -5.77 | 7.88E-07 | *** |
| Age | -0.001 | 0.003 | -0.33 | 0.740919 | |
| mFD | 0.154 | 0.311 | 0.49 | 0.623568 | |

Residual standard error: 0.1308 on 43 degrees of freedom
Multiple R-squared: 0.4368, Adjusted R-squared: 0.3975
F-statistic: 11.12 on 3 and 43 DF, p-value: 1.576e-05

**Table 6**

**rpInsula-putamen = 1 + FTND + age + mFD**

| | Estimate | Std.Error | t-value | Pr(>\|t\|) | |
|---|---|---|---|---|---|
| (Intercept) | 0.376 | 0.095 | 3.94 | 0.000294 | *** |
| FTND | -0.044 | 0.009 | -5.21 | 5.17E-06 | *** |
| Age | -0.001 | 0.002 | -0.59 | 0.558069 | |
| mFD | -0.024 | 0.273 | -0.09 | 0.928925 | |

Residual standard error: 0.1149 on 43 degrees of freedom
Multiple R-squared: 0.3907, Adjusted R-squared: 0.3482
F-statistic: 9.193 on 3 and 43 DF, p-value: 8.122e-05

**Table 7**

**lpInsula-putamen = 1 + FTND135 + F1246 + age + mFD**

| | Estimate | Std.Error | t-value | Pr(>\|t\|) | |
|---|---|---|---|---|---|
| (Intercept) | 0.267 | 0.092 | 2.90 | 0.00596 | ** |
| F135 | 0.013 | 0.013 | 1.00 | 0.32094 | |
| F246 | -0.066 | 0.010 | -6.32 | 1.37E-07 | *** |
| Age | -0.003 | 0.002 | -1.30 | 0.20174 | |
| mFD | 0.182 | 0.273 | 0.67 | 0.50994 | |

Residual standard error: 0.1149 on 42 degrees of freedom
Multiple R-squared: 0.5267, Adjusted R-squared: 0.4816
F-statistic: 11.68 on 4 and 42 DF, p-value: 1.817e-06

**Table 8**

**rpInsula-putamen = 1 + FTND135 + F1246 + age + mFD**

| | Estimate | Std.Error | t-value | Pr(>\|t\|) | |
|---|---|---|---|---|---|
| (Intercept) | 0.226 | 0.083 | 2.72 | 0.0095 | ** |
| F135 | 0.011 | 0.012 | 0.93 | 0.3574 | |
| F246 | -0.052 | 0.009 | -5.51 | 1.98E-06 | *** |
| Age | -0.002 | 0.002 | -1.01 | 0.3197 | |
| mFD | 0.074 | 0.246 | 0.30 | 0.7665 | |

Residual standard error: 0.1035 on 42 degrees of freedom
Multiple R-squared: 0.4554, Adjusted R-squared: 0.4035
F-statistic: 8.78 on 4 and 42 DF, p-value: 3.032e-05

**Table 9**

**lvaInsula-precuneus = 1 + FTND135 + F1246 + age + mFD**

| | Estimate | Std.Error | t-value | Pr(>\|t\|) | |
|---|---|---|---|---|---|
| (Intercept) | -0.028 | 0.102 | -0.27 | 0.78951 | |
| F135 | -0.033 | 0.014 | -2.36 | 0.02318 | * |
| F246 | -0.036 | 0.012 | -3.05 | 0.00397 | ** |
| Age | 0.001 | 0.003 | 0.38 | 0.70913 | |
| mFD | 0.276 | 0.304 | 0.91 | 0.36884 | |

Residual standard error: 0.1278 on 42 degrees of freedom
Multiple R-squared: 0.4195, Adjusted R-squared: 0.3642
F-statistic: 7.589 on 4 and 42 DF, p-value: 0.0001074

**Table 10**

**ldaInsula-precuneus = 1 + FTND135 + F1246 + age + mFD**

| | Estimate | Std.Error | t-value | Pr(>\|t\|) | |
|---|---|---|---|---|---|
| (Intercept) | 0.213 | 0.121 | 1.75 | 0.0868 | ^ |
| F135 | -0.034 | 0.017 | -2.03 | 0.0488 | * |
| F246 | -0.036 | 0.014 | -2.63 | 0.0119 | * |
| Age | -0.002 | 0.003 | -0.69 | 0.4914 | |
| mFD | 0.237 | 0.360 | 0.66 | 0.5148 | |

Residual standard error: 0.1513 on 42 degrees of freedom
Multiple R-squared: 0.3491, Adjusted R-squared: 0.2871
F-statistic: 5.632 on 4 and 42 DF, p-value: 0.001009

**Table 11**

**rvaInsula-precuneus = 1 + FTND135 + F1246 + age + mFD**

| | Estimate | Std.Error | t-value | Pr(>\|t\|) | |
|---|---|---|---|---|---|
| (Intercept) | -0.157 | 0.134 | -1.17 | 0.2474 | |
| F135 | -0.042 | 0.019 | -2.27 | 0.0281 | * |
| F246 | -0.027 | 0.015 | -1.74 | 0.0898 | ^ |
| Age | 0.003 | 0.003 | 0.72 | 0.4733 | |
| mFD | 0.805 | 0.399 | 2.02 | 0.0497 | * |

Residual standard error: 0.1674 on 42 degrees of freedom
Multiple R-squared: 0.3209, Adjusted R-squared: 0.2563
F-statistic: 4.963 on 4 and 42 DF, p-value: 0.002284

**Table 12**

**rdaInsula-precuneus = 1 + FTND135 + F1246 + age + mFD**

| | Estimate | Std.Error | t-value | Pr(>\|t\|) | |
|---|---|---|---|---|---|
| (Intercept) | 0.055 | 0.099 | 0.56 | 0.58192 | |
| F135 | -0.046 | 0.014 | -3.33 | 0.00184 | ** |
| F246 | -0.024 | 0.011 | -2.14 | 0.03854 | * |
| Age | 0.000 | 0.003 | -0.15 | 0.88365 | |
| mFD | 0.441 | 0.294 | 1.50 | 0.1415 | |

Residual standard error: 0.1236 on 42 degrees of freedom
Multiple R-squared: 0.4304, Adjusted R-squared: 0.3762
F-statistic: 7.934 on 4 and 42 DF, p-value: 7.386e-05

Significance codes: ***0.001 **0.01 *0.05 ^0.1

**Table 13**

**FTND exploratory factor analyses comparison**

| | | Current study | | Haddock et al, 1999 | |
|---|---|---|---|---|---|
| **FTND items** | **Question** | **DS** | **MS** | **DS** | **MS** |
| **FTND1** | How soon after you wake up do you smoke your first cigarette? | 0.57 | 0.43 | 0.68 | 0.43 |
| **FTND2** | Do you find it difficult to keep from smoking in places where it is not allowed? | 0.54 | 0.17 | 0.63 | -0.1 |
| **FTND3** | Which cigarette would you hate most to give up? | -0.03 | 0.89 | 0.04 | 0.79 |
| **FTND4** | On average, how many cigarettes are you currently smoking each day? | 0.72 | -0.18 | 0.76 | 0.12 |
| **FTND5** | Do you smoke more frequently during the first hours after waking than during the rest of the day? | 0.06 | 0.73 | 0.11 | 0.73 |
| **FTND6** | Do you smoke if you are so ill that you are in bed most of the day? | 0.73 | -0.02 | 0.74 | 0.14 |
| | | Principal components analysis followed by Varimax rotation explaining 54% of variance (N=47). | | Principal components analysis followed by Varimax rotation explaining 51.5% of variance (N=4042). | |
